# Formation of correlated chromatin domains at nanoscale dynamic resolution during transcription

**DOI:** 10.1101/230789

**Authors:** Haitham A. Shaban, Roman Barth, Kerstin Bystricky

## Abstract

Intrinsic dynamics of chromatin contribute to gene regulation. How chromatin mobility responds to genomic processes and whether this response relies on coordinated movement is still unclear. Here, we introduce an approach called Dense Flow reConstruction and Correlation (DFCC) to quantify correlation of chromatin motion with sub-pixel sensitivity at the level of the whole nucleus. DFCC is based on reconstructing dense global flow fields of fluorescent images acquired in real-time. By simulating variations in microscopic and dynamic parameters, we demonstrate that our approach is robust and more accurate than other methods to estimate flow fields and spatial correlations of dense structures such as chromatin. We applied our approach to analyze stochastic movements of DNA and histones based on direction and magnitude at different time lags in human cells. We observe long-range correlations extending over several μm between coherently moving regions over the entire nucleus. Spatial correlation of global chromatin dynamics was reduced by inhibiting elongation by RNA polymerase II and abolished in quiescent cells. Furthermore, quantification of spatial smoothness over time intervals up to 30 seconds points to clear-cut boundaries between distinct regions, while smooth transitions in small (<1 μm) neighborhoods dominate for short time intervals. Clear transitions between regions of coherent motion indicate directed squeezing or stretching of chromatin boundaries suggestive of changes in local concentrations of actors regulating gene expression. The DFCC approach hence allows characterizing stochastically forming domains of specific nuclear activity.

**Significance Statement:** Control of gene expression relies on modifications of chromatin structure and activity of the transcription machinery. However, how chromatin responds dynamically to this genomic process and whether this response is coordinated in space is still unclear. We introduce a novel approach called Dense Flow reConstruction and Correlation (DFCC) to characterize spatially correlated dynamics of chromatin in living cells at nanoscale resolution. DFCC allows us to detect chromatin domains in living cells with long range correlations over the entire nucleus. Furthermore, transitions between domains can be quantified by the newly introduced smoothness parameter of local chromatin motion. The DFCC approach permits characterizing stochastically forming domains of other DNA dependent activity in any cell type in real time imaging.

## Introduction

The spatial organization of chromosomes is characterized by short- and long-range contacts bringing chromosome domains into spatial proximity and creating chromosome territories (CT) detectable by fluorescent in situ hybridization (FISH) (1, 2) and high-throughput chromosome conformation capture (Hi-C) (3) in human cells. Live cell imaging of CTs identified random and directed motion of subchromosomal foci and suggested similarities in dynamic behavior between distinct CTs (4). On the time scale of several seconds, chromatin was shown to move coherently irrespective of CT boundaries implying a transient mechanical coupling between chromatin over a few microns (5, 6). Fluctuations in chromatin architecture occur over a large range of spatio-temporal scales during regulatory processes (3, 7, 8). This range of variations makes studying of molecular organization and dynamic processes of the whole genome challenging. In particular, actively transcribed genes depend on chromatin dynamics to fine-tune expression levels (9). Active genes sometimes gather dynamically to share the same transcription sites (10, 11) within which the range of chromatin movement is thought to correlate with mRNA production and enhancer activity (10). Coordinated relocalisation or extrusion of activated genes to the surface of CTs is coherent with the idea that this process allows reaching shared transcription factors (12–14). The mechanism of such movements is unknown.

Chromatin motion during processes related to genome function in mammalian cells can be studied by fluorescence live imaging. Most of these studies rely on labeling single loci or arrays of repeated DNA sequences with assistance of gene editing techniques (15–20). Single particle tracking demonstrated that motion of the tagged DNA loci in cells is sub-diffusive although super-diffusive behavior was reported (18, 19). The heterogeneity in motion of telomeres is particularly striking (21, 22). In agreement, large fluctuations in sub-diffusive behavior were also determined for H2B-GFP imaged with sub-second time intervals (23, 24). To gain a true understanding of the physical nature of a long fiber structure such as genomic chromatin and how this fiber behaves on a global scale, chromatin has to be analyzed at a large scale across the entire nucleus. Recently, a dynamic analysis based on correlation spectroscopy of time-resolved imaging using particle imaging velocimetry (PIV) (6) reported a global view on the chromatin motions. However, a relatively large interrogation window size of more than one micrometer was set to estimate the displacement vectors. Dynamic changes within the set window cannot be considered, hence missing the contribution of genomic processes to local chromatin motion.

Here, we introduce an approach called Dense Flow reConstruction and Correlation (DFCC) to quantify the correlation of chromatin motion with sub-pixel sensitivity at the level of the whole nucleus. DFCC provides sub-diffraction vectorial information, based on reconstructed dense global flow fields of a series of diffraction limited fluorescent images. The sample pixel size defines the dynamic resolution of the results independently of its dimensions (here down to 65 nm). We use Optical Flow (OF) to estimate the direction and amplitude of the motion of fluorescent labeled DNA and histones over a 30 second time interval at 5 fps, and confirm that an OF formulation is more sensitive than PIV for studying the motion of intracellular objects (25). We calculate the spatial and temporal correlation based on both direction and amplitude of each displacement vector, and quantify characteristic length scales of correlated motion with nano-scale resolution. Estimation of the smoothness of flow fields across the whole nucleus at different transcriptional stages reveals coherently moving chromatin domains.

## Results and discussion

### Comparison of Optical Flow methods for precise estimation of chromatin dynamics by simulations

In order to evaluate the accuracy of different OF methods quantitatively for chromatin motion, ground truth data is required, but unavailable for biological systems. To overcome this lack we simulated images recapitulating our experimental conditions with a range of microscopic (labeling density, Signal-to-Noise ratio (SNR)) and dynamic (diffusion coefficient, number of independently moving domains) parameters. In this study, we considered five Optical Flow methods that cover four types of OF, namely differential methods (Horn and Schunck (HS); Lucas and Kanade (LK) formulations) (26, 27), region-based matching (hereafter Particle Image Velocimetry – PIV) (28), phase-based methods (29) and SIFT-based methods (30). Details of all the tested algorithms can be found in Supplementary Note 1. We evaluated the performance of the different methods by determining the angular error (AE) and the endpoint error (EE) in simulated data sets (Materials and Methods). The simulation of data samples was carried out by randomly placing emitters with a defined density in a given volume with varying SNR as described in (Supplementary Note 1, Supplementary Figure S1). A series of two images was simulated where emitters undergo Brownian diffusion and therefore are displaced from one image to another. We simulated a density ranging from 0.02 to 2.5 emitters per pixel representing the spatial variation of chromatin compactness within hetero- and euchromatin (23) (Supplementary FigureS1a-c). Further, we varied the particles’ diffusion coefficient as well as the number of coherently moving domains which could potentially reflect chromatin motion at different length scales (Supplementary FigureS1f-g). The images were then subjected to OF algorithms in order to reconstruct the direction and magnitude of the emitters’ movements. Regions within the emitters are forced to undergo coherent motion were superimposed as described in (Supplementary Note 1).

First we tested the impact of labeling density on the accuracy of the methods to define vectors’ AE and EE. We noticed that by increasing labeling density, error measures increased for all methods. A jump was seen in the EE of the HS formulation once the density approaches~1/*px*. Our simulations showed that, even with labeling density increasing 150-fold, the AE maximally increased 3.9-fold and EE increased 3.4-fold (Figure 1). The tested methods therefore did not scale linearly with increasing density but were robust with respect to varying emitter densities. Although HS and PIV (window size 16×16 pixels) performed best, HS had the lowest AE (<10 deg), 2-fold better than PIV (16×16 pixels).

**Figure 1:**
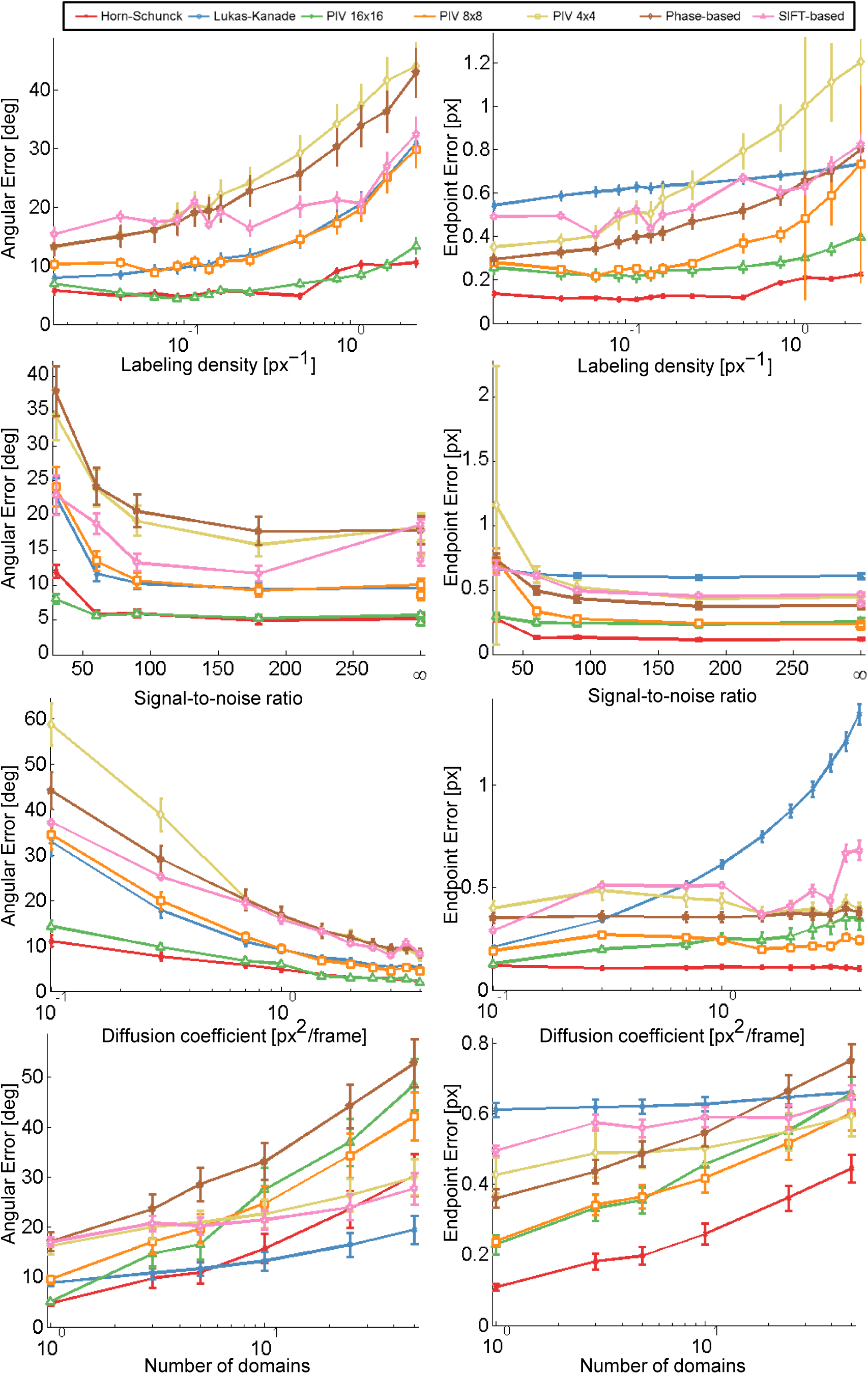
Performance of different Optical Flow methods. Angular and endpoint error for each method is shown under variation of static (labeling density and Signal-to-Noise ratio) and dynamic parameters (diffusion coefficient and number of domains). Angular error is shown in the left column, endpoint error in the right; parameters are shown in rows and vary from low to high values. The response of each method in terms of AE and EE is plotted (see Supplementary Table 1 and Supplementary Note 1). Error bars are symmetric and correspond to the standard deviation from 10 simulations. Lower AE and EE mean more accurate estimates.

Next, we considered variations in SNR due to characteristics of the specimen and intrinsic signal-dependent Poisson noise (Supplementary Figure S1d-e). Figure 1 shows that the accuracy of all methods decreases with noise and a characteristic step at *SNR* ≈ 90 can be observed. The EE of PIV methods using a small window size (8x8 pixels or smaller) was particularly sensitive to a low SNR because cross-correlation methods strongly respond to noise. When the noise levels are high, a displacement peak may become smaller than surrounding noise peaks and the probability increases that an erroneous noise peak is chosen (31). Especially for small displacements, this results in a bias towards larger displacements and therefore an increase in the EE.

Optical Flow methods should be sensitive to displacement magnitude in order to capture the temporal and spatial amplitude of chromatin dynamics, typically characterized by diffusion coefficients ranging from10^−2^ to10^−3^ μm^2^/s (6, 23). Increasing the emitters’ diffusion coefficient, i.e. increasing the distance by which each emitter is allowed to move per time step, enhanced the accuracy in determining the direction of displacement. OF methods using coarse-to-fine estimation schemes did not show substantially different trends than methods without a pyramidal structure (Figure 1). For the SIFT-based method, the magnitude of estimated vectors is unrestricted, and therefore displacements investigated in this study did not impose limitations on the method. However, the EE of the LK formulation was substantially less robust against variations in the displacement’s magnitude than other formulations.

Varying the number of independently moving domains showed that errors increased with greater complexity in motion independently of the method employed. For 10 domains and more, the LK-based method outperformed the HS formulation in terms of AE. Nevertheless, if the accuracy of the method is assessed by EE, the HS method performed substantially better than all other methods. Note that results from the SIFT-based method were the most stable to variations in imaging conditions among all investigated methods. The local LK formulation displayed relatively a high EE for all simulation parameters. The formulation requires a Tikhonov regularization (32), which resulted in a bias towards smaller displacements and therefore increased the EE (Supplementary Note 1).

Several PIV methods differing by their interrogation window size were tested. The accuracy achieved by these methods was dependent on the window size. Although the AE of PIV using a window size of 16x16 pixels was as low as the one using the HS method, PIV failed to identify different domains of coherent motion. Based on our simulations, we determined that the interrogation window size used in PIV should be equal or larger than the expected maximal displacement. Otherwise, emitters moving out of the interrogation window create errors resulting from calculations of cross-correlation from signal loss. In the present study, the maximum distance was 4 px/frame, and therefore even PIV using the smallest window size of 4×4 pixels was capable to estimate direction of motion. Since the calculation of the cross-correlation largely depended on window size, the window-size must be carefully adjusted to the data. Multiple small independently moving objects within the same subregion might lead to erroneous results due to several independent motions within the pattern. Careful adjustment of the window size is particularly difficult when studying chromatin dynamics (6), where the expected density of emitters and their dynamic behavior is hard to predict.

On average, the HS-based method outperformed all other methods and was thus chosen as the most reliable for the experimental analysis hereafter. In conclusion, our simulations show that the HS based OF method is appropriate for studying chromatin dynamics based on fluorescence imaging.

### Correlation modeling of dense motion fields

Vector fields produced by Optical Flow algorithms may be considered as two independent scalar fields representing direction and magnitude of the vector field. Each value in these fields can be described as a stochastic variable which led us to consider the scalar fields as random fields. This allows parametrization of the correlation of chromatin dynamics in each random field using the Whittle-Màtern (WM) model (33, 34):

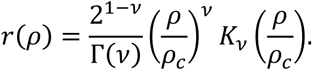

Where, Γ(·) is the gamma function; *K_v_*(·) is the modified Bessel function of the second type of order *v*, *ρ_c_* is the correlation length, and *v* is a smoothness parameter (Supplementary Note 2). The WM model has important advantages for modeling spatial processes by including a parameter which characterizes the smoothness of the corresponding random fields. Large *v* means that the underlying spatial process is smooth in space, whereas the process is considered as rough for small *v* (35, 36) (Supplementary Note 2). An analogy between the smoothness of a random field and its differentiability can be drawn (provided *v* ≥ 1) (37). The association of the smoothness parameter to the existence of directional derivatives gives rise to the concept that a smooth field does not exhibit singular points and is continuous in the domain of computation. Smooth fields are uncommon for natural processes, but the parametrization of smoothness allows identifying sharp transitions (e.g. object boundaries, singular points) in a quantified manner.

### Dense flow correlation of chromatin dynamics based on real time imaging in 2D

We determined flow fields of chromatin motion based on real time imaging (a series of 150 single plane images acquired at 5 fps) of a single U2OS nucleus expressing H2B-GFP (Figure 2a-b) and compared the response of the five methods under consideration. The empirical correlation was calculated and projected onto the one-dimensional space lag *ρ* from two successive representative images (see Materials and Methods, Figure 2 c-d). We applied the WM model to extract the correlation length and smoothness parameter (Figure 2e). Figure 3 shows the mapped direction and magnitude of each single vector across a single entire nucleus (Figure 3a-b). In the flow field of the LK method, one could hardly observe large regions of coherent motion, both in direction and magnitude. The flow field was rough, showing many small areas and therefore correlation dropped right after the zero space lag (Figure 3c). This behavior was seen in both direction and magnitude. Consistent with our simulations, the window size of PIV methods influenced their ability to distinguish between regions of coherent flow. PIV with a 16x16 pixel window size showed numerous small regions of coherent motion (in direction only) and did not show clear distinction between direction and magnitude of spatial correlation. Moreover, correlation dropped sharply after the zero space lag for both direction and magnitude. The fact that small regions arose by using the 16×16 pixel window size may be due to several independent motion areas within the interrogation window. Shifting the window from one pixel to another and including intensity information further apart from the vector to be estimated may therefore substantially influence vector estimation, even for adjacent pixels. In other words, averaging a vector over more than a micrometer for chromatin structures introduced errors in both direction and magnitude of the detected motion, and therefore led to inaccurate estimation of motion and correlation as shown by simulating a high number of independently moving areas. The SIFT method showed great similarity between the correlation of direction and magnitude, which is likely due to vector quantization (integer values for the x and y-components are allowed only, see Supplementary Note 2). Only the HS based method was able to identify changes in correlation over several space lags, and to distinguish between direction and magnitude.

**Figure 2:**
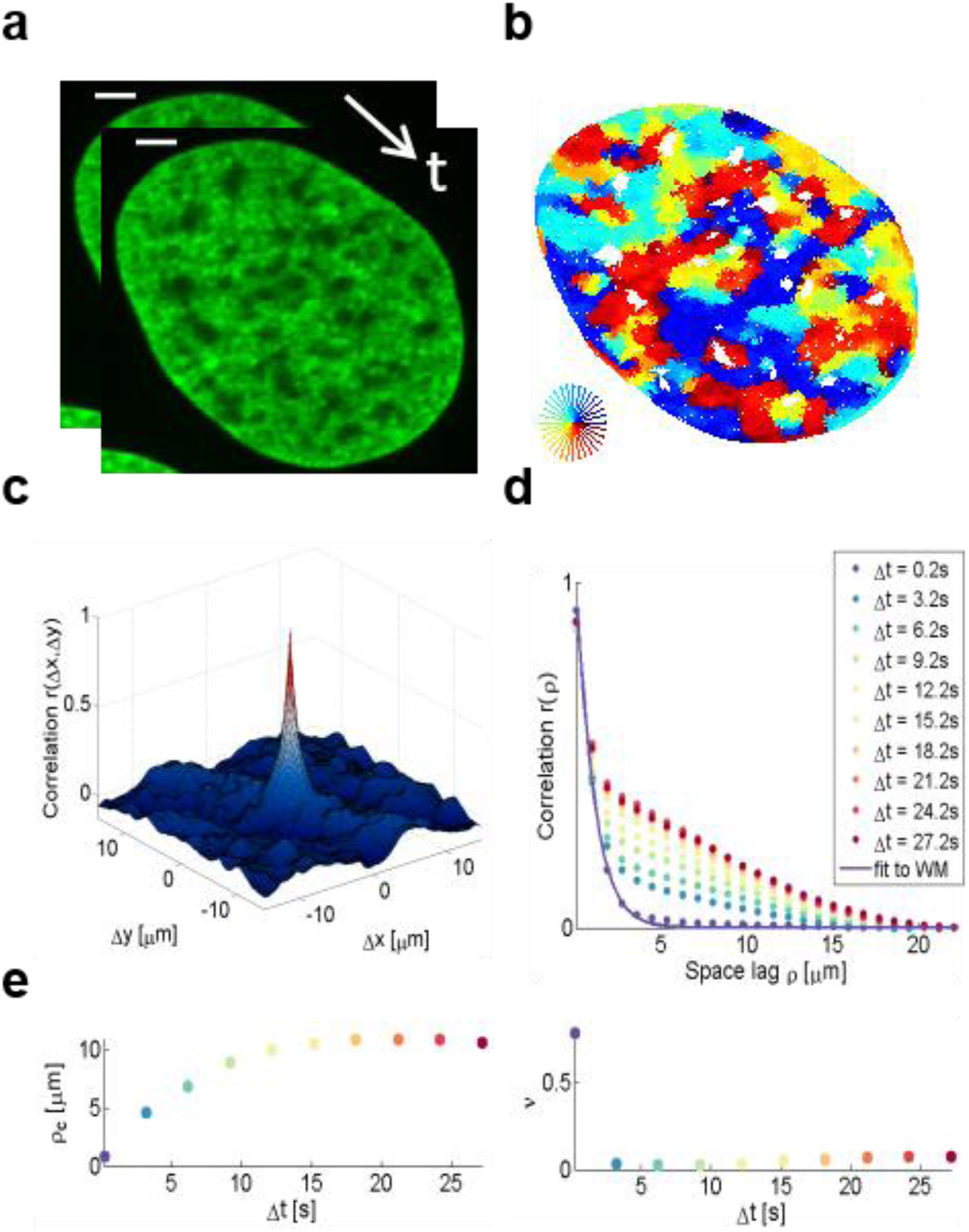
Schematic representation of the correlation analysis. **a)** Two microscopy images (U2OS cells expressing H2B-GFP) were acquired with temporal resolution 200 ms. Scale bar is 3 μm. **b)** The flow field between the input images was estimated by the Horn-Schunck formulation and color-coded as indicated in the lower right. Pixels in nucleoli appear empty due to the lack of intensity information. **c)** Correlation calculation in two dimensions. **d)** Empirical correlation (example shown for direction) was calculated as a function of space lag and fitted to the Whittle-Màtern correlation model. Flow fields were estimated at every accessible time lags within the image series and each of them was fitted (example time lags shown only). **d)** Correlation length *ρ_c_* and the smoothness parameter *v* were derived from the regression and shown over the time lag. The parameters were averaged for each time interval over all accessible time points. Note that with increasing time interval, less time points are available and therefore, the standard deviation (not shown) increases.

**Figure 3:**
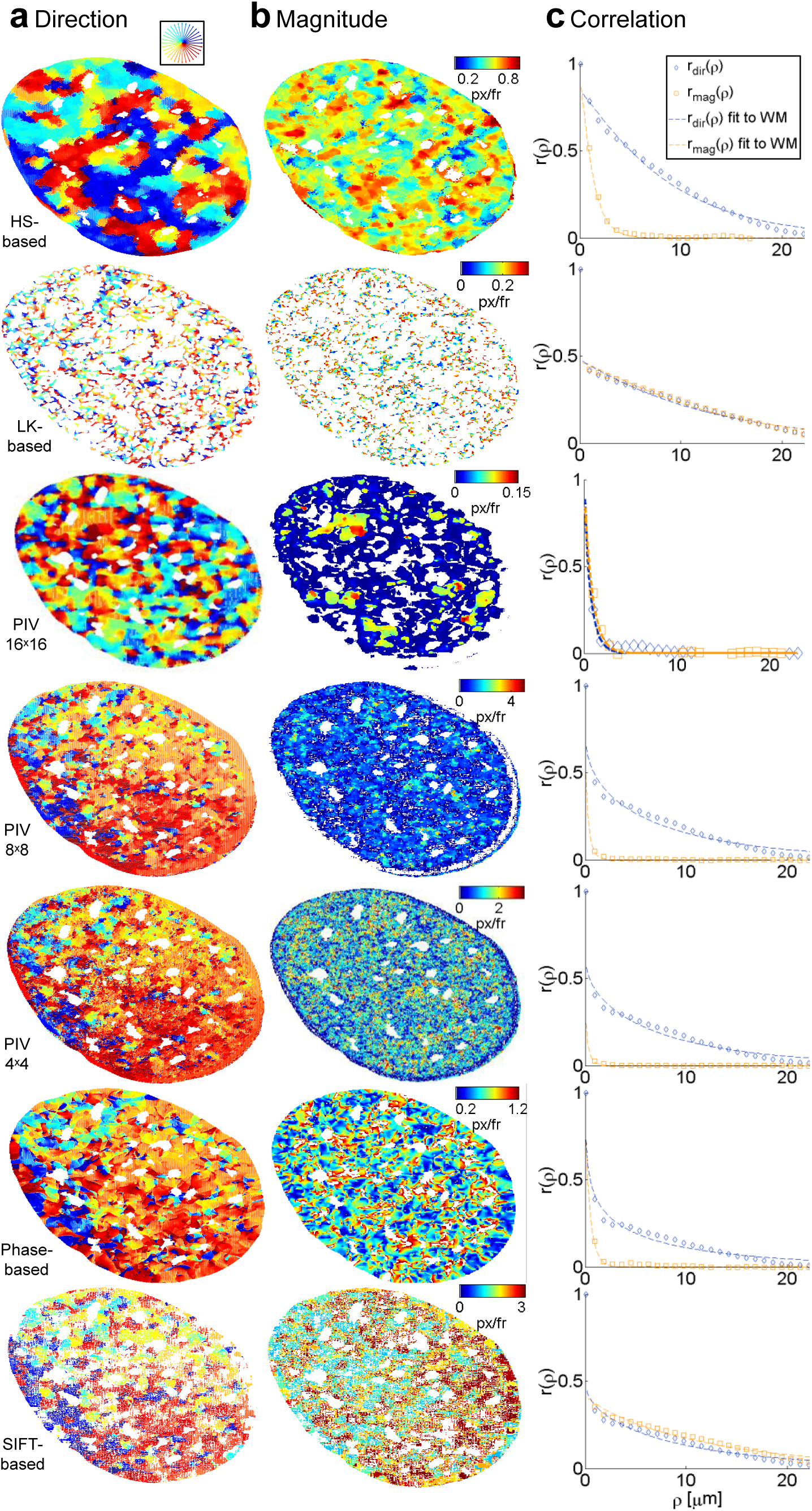
Representative visualization of flow fields for the analysis of a single U2OS cell expressing H2B-GFP. Rows correspond to the different investigated methods. **a)** Flow fields are color-coded by the direction of displacement vectors. **b)** Magnitude of the corresponding flow vector color-coded from low to high as indicated by color bars. **c)** Empirical correlation for direction (blue) and magnitude (orange) and corresponding fits to the Whittle-Màtern model (solid lines) over space lags.

Finally, the Whittle-Màtern model enabled quantification of the smoothness parameter. We illustrate the behavior of the smoothness parameter by means of estimated flow fields using HS and SIFT-based methods (Supplementary Note 2, Supplementary Figure S3). The variance of the horizontal and vertical gradient (directional) fields was calculated (Supplementary Figure S3b-c). Large gradients correspond to sharp transitions (recall e.g. the gradient of a step function) and small gradients correspond to domains separated by smooth transitions. Therefore, a decrease in the variance of the gradient fields corresponds to an increase of the smoothness parameter. The HS based method showed few large areas of coherent motion. Inside these regions, the field varied only slightly and the gradient within these areas was small. At motion boundaries, however, direction of vectors changed abruptly (Supplementary Figure S3b-c). This abrupt change in direction is likely due to changes in the chromatin surroundings and interactions, as for example those brought about by factors involved in gene expression.

### Chromatin dynamics at nano-scale resolution reveals transcription dependent long range correlation

We applied DFCC to analyze movements of chromatin in the entire nucleus and to assess whether chromatin dynamics were sensitive to transcriptional state. Correlation of coherent chromatin motion was calculated using two different fluorescent labels: DNA labeled SiR–Hoechst and H2B fused to GFP in human osteosarcoma U2OS cells. Images were recorded in cells grown in medium containing serum (actively transcribing state) and in cells starved in serum for 24H (inactive state). Analysis of chromatin motion in cells grown in medium containing serum showed that the correlation length for both DNA and H2B was time dependent, reaching a maximum correlation length (*ρ_c_* ≈ 11μ*m*) at 18.2s (Figures 4c-d, Figure 5c-d). Correlation over long range is consistent with the notion that active transcription occurs in numerous regions of decompacted, open chromatin (38). Interestingly, some regions of coherent motion comprise smaller areas of the scale of a few hundred nanometers within them, whose patterns deviate slightly from the direction of the bulk of the coherent domains (enlarged areas, Figures 4b and 5b) suggesting that territories of a single chromosome are not necessarily moving in the same direction or manner. This result concurs with previous observations that coherent regions of H2B-GFP movement spanned across several dNTP labeled CTs (6). Furthermore, flow fields comprised distinct regions of vortex-like motion of a few hundred nanometers length scale (Figure 6a, Supplementary Figure 6a-b), which might support the hypothesis that genes are moving to share transcription factors within transcription factories or hubs (10, 11). The resolution afforded by our approach permits analysis of sub-micron motion within domains which may represent hubs of active DNA-associated processes.

**Figure 4:**
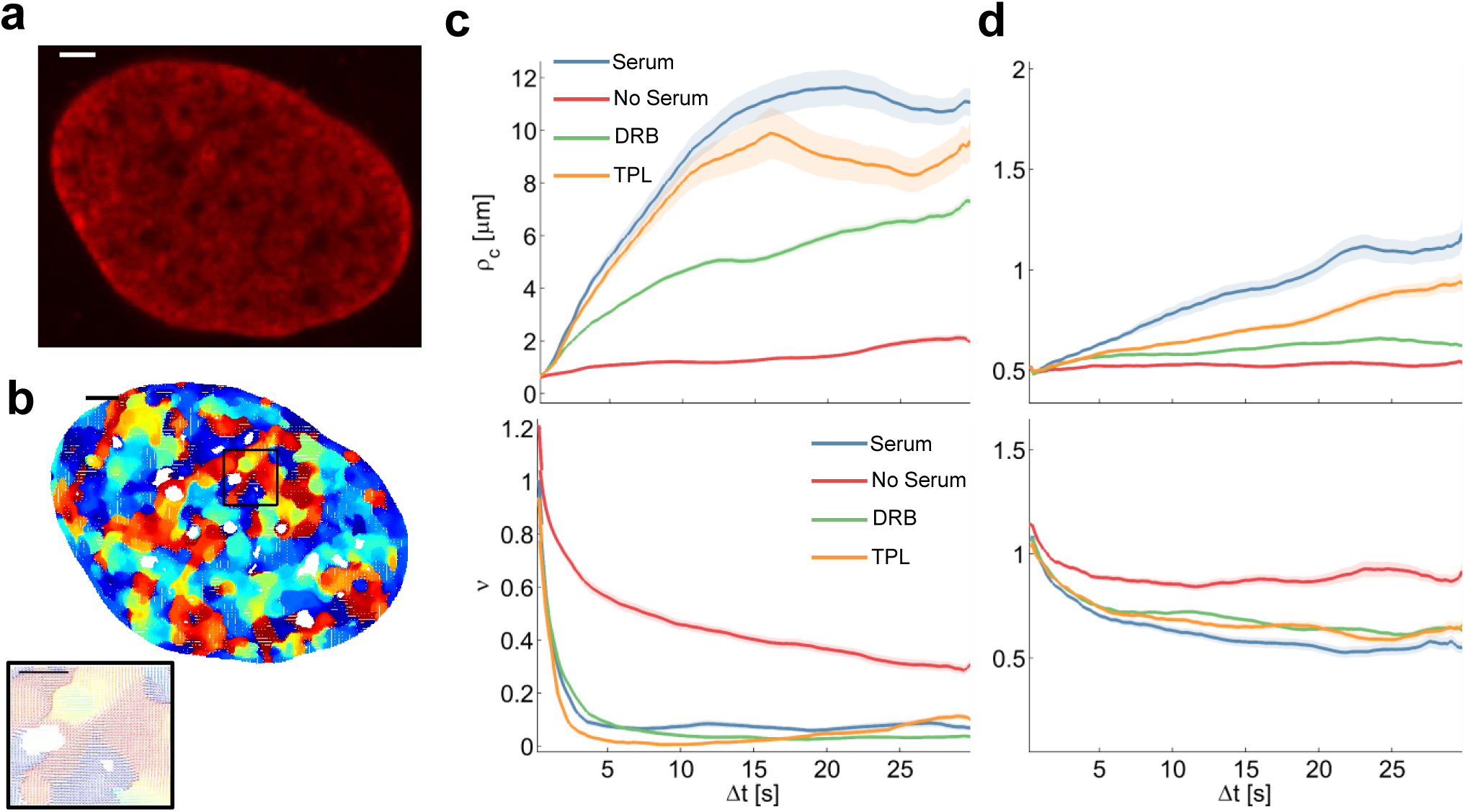
Correlation length and smoothness in direction and magnitude of DNA in U2OS cells using the Whittle-Màtern model **a)** A fluorescence microscopy image of a nucleus where DNA was labeled using Sir-Hoechst; scale bar is 3 μm. **b)** Flow field for Δ*t* = 0.2 *s* and enlarged region (*right*) of the black rectangle; the field is color-coded according to the direction of the displacement. Scale bar is 3 μm (*left*) and 1 μm (*right*). **c)** Correlation length (*top*) and smoothness parameter (*bottom*) calculated from regression of empirical correlation functions over time for directional correlation of flow fields. Different colors correspond to different conditions. Shaded error bars correspond to the standard deviation over 18 nuclei per condition. **d)** As **c)** for the vectors’ magnitude.

**Figure 5:**
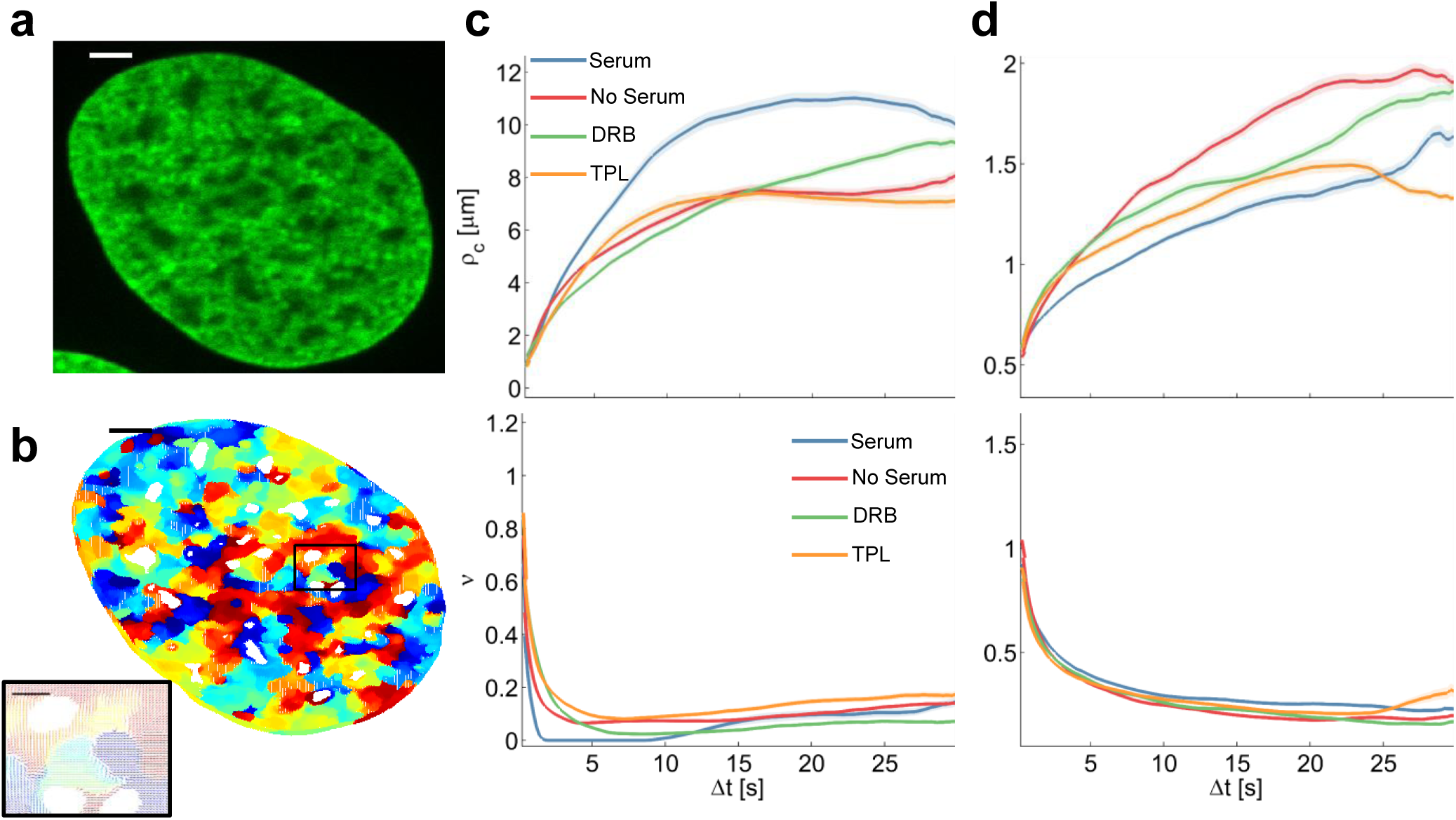
Correlation length and smoothness in direction and magnitude of H2B-tagged GFP in U2OS cells using the Whittle-Màtern model. **a)** A fluorescence microscopy image of a nucleus expressing H2B-GFP; scale bar is 3 μm. **b)** Flow field for Δ*t* = 0.2 *s* and zoomed-in region *(right)* of the black rectangle; the field is color-coded according to the direction of the displacement. Scale bar is 3 μm (*left*) and 1 μm (*right*). **c)** Correlation length (*top*) and smoothness parameter (*bottom*) calculated from regression of empirical correlation functions over time lag for directional correlation of flow fields. Different colors correspond to different conditions. Shaded error bars correspond to the standard deviation over 19 nuclei per condition. **d)** As **c)** for the vectors’ magnitude.

**Figure 6:**
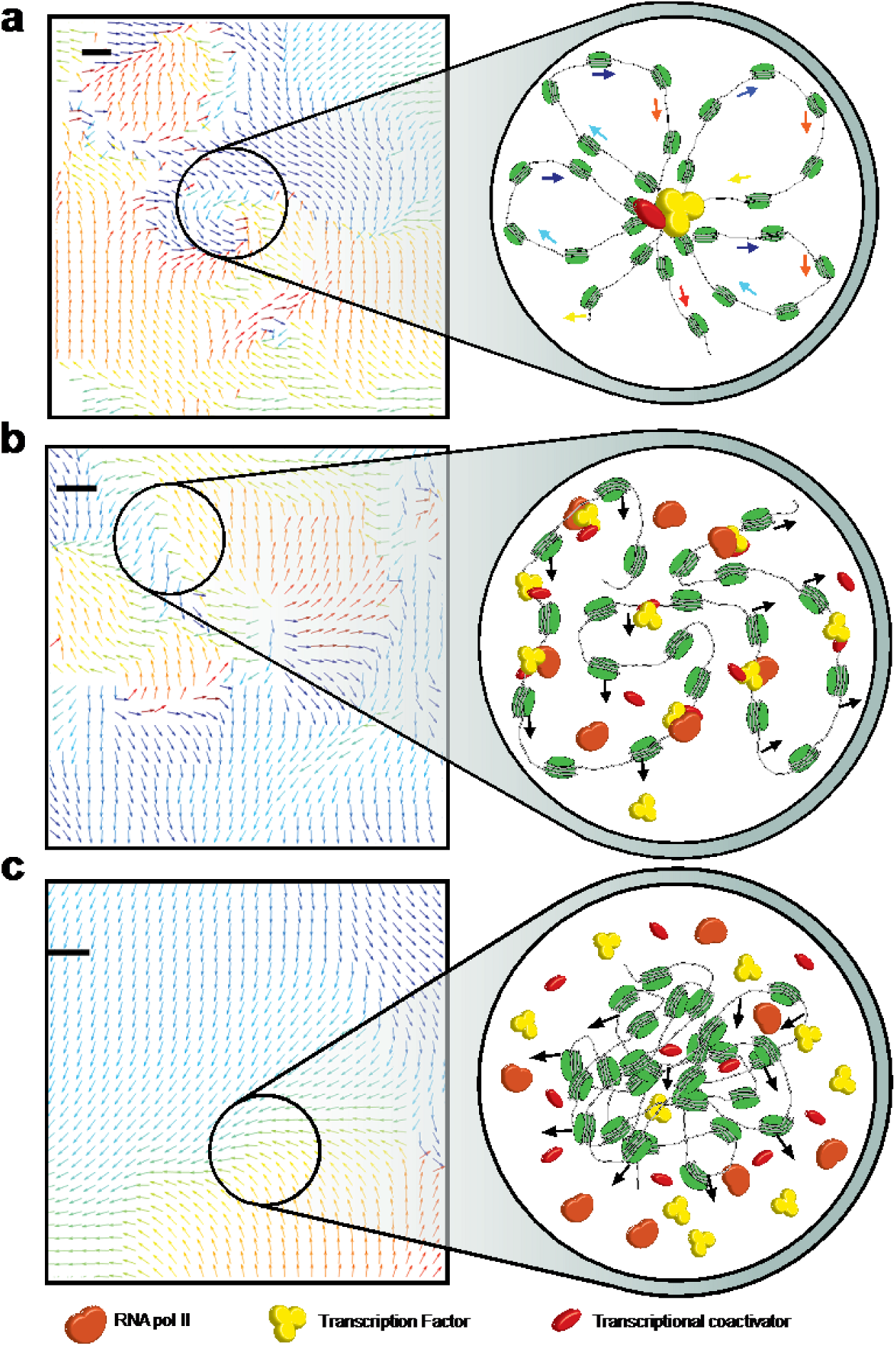
Representative magnification of flow fields and models of proposed mechanisms. **a)** An example region of converging flow (vortex) observed in the mean direction over 30 s is enlarged (Supplementary Figure S4). In- and outward directed motion may be due to loop formation during the transcription cycle as illustrated. **b)** Visual change in flow smoothness. The observed change in smoothness between serum stimulation (b) and starvation **(c)** in case of DNA probing is visualized by representative regions (Supplementary Figure 4). The observed low smoothness in case of serum stimulation is due to chromatin decompaction as illustrated. Chromatin is able to move rather freely and RNA pol II as well as transcription factors can bind to DNA. Coherent motion and sharp motion boundaries were observed and indicate that local chromatin regions converge towards shared transcription factors. **d)** In case of serum starvation, correlation drops and coherently moving regions seamlessly interrelate. An increase in chromatin compaction causes DNA-DNA interactions to occur more frequently which causes spatial transitions in directional chromatin motion to be smooth. Furthermore, chromatin compaction has an inhibiting effect on protein dynamics and protein binding to DNA is therefore hampered. Scale bars are 200 nm.

Quantification of spatial smoothness over large time intervals points to clear-cut boundaries between distinct regions, while smooth transitions in small neighborhoods dominate for short time intervals. These transitions between regions of distinct motion indicate directed squeezing or stretching of chromatin boundaries suggestive of changes in local concentrations of actors regulating gene expression (Figure 6b, Supplementary Figure 6c).

In contrast, chromatin motion occurred in numerous small domains with smooth transitions in serum starved cells over 30 seconds (Figure 4c-d). The spatial correlation (both direction and magnitude) of SiR-labelled DNA showed almost no time dependence and fitting by the MW model yielded a correlation length of less than one micrometer (Figure 4c-d). The nearly constant spatial correlation is due to the fact that serum-starved cells are in quiescence, in which chromatin fibers are more compact (38, 39). In conclusion, short correlation length and comparably high smoothness of flow may be due to condensed chromatin regions (Figure 6c, Supplementary Figure 6d).

However, motion of H2B-GFP in starved cells yielded a maximum directional correlation length of *ρ_c_* ≈ 7 μ*m*, half the length calculated in cells with serum stimulation (Figure 5c-d), but greater than for motion of SiR-labelled DNA. Differences in the amplitude of correlation between SiR-labelled DNA and H2B-GFP might be due to variations in the labeling density affecting determination of domains and consequently correlation length calculations. In other words, only 10-20 % of chromatin contains labelled H2B-GFP (40) may represent the least compacted fraction of total chromatin characterized by greater correlation length and rougher of flow. The correlation in serum starved cells could stem from free H2B-GFP molecules not incorporated into chromatin.

### Coordinated movements within domains correlate with RNA polymerase II activity

We further assessed the effect of transcription inhibitors on initiation or elongation of transcription by pretreating the cells with 5,6-Dichloro-1-β-D-ribofuranosylbenzimidazole (DRB) and Triptolide (TPL), respectively. Inhibition of RNA polymerase II (RNA pol II) had greater and more diverse effects imaging SiR-labelled DNA than H2B-GFP (Figure 4c-d). Correlation length and magnitude of DNA motion was reduced in cells treated with DRB and only slightly affected by TPL compared to cells grown in serum, suggesting that elongation maintains dynamic transitions (Figure 4c-d). The relative smoothness of motion, however, remained similar to the one determined when cells were grown in serum, demonstrating that rough boundaries correspond to the presence of RNA pol II and cofactors in certain domains of the nucleus. Correlation length and relative smoothness of the motion of H2B-GFP upon RNA pol II inhibition were indistinguishable from the values determined in serum starved cells reinforcing the idea that active transcription drives H2B-GFP nuclear domain formation (Figure 5c-d). The magnitude of correlation increased at longer time intervals in inhibitors treated nuclei but did not reach values determined in serum starved conditions. Hence, the remaining association of polymerases and cofactors during initiation (DRB causes RNA pol II to detach within the first exon (41)) or remaining elongation (TPL precludes RNA pol II from binding to the promoter but RNA pol II still elongating after 15 min treatment will proceed (42)) and possibly the presence of nascent mRNA influences dynamic behavior of chromatin.

The clear difference of coordinated movements between labelled DNA and H2B-GFP may also be exacerbated by loss of histones resulting from starvation- or inhibitor-induced stress (43). Our results showed similarities between actively transcribed (+serum) and initiation halted chromatin, where long scale correlation was seemingly due to chromatin compaction. Given that neither TPL nor DRB block RNA polymerase activity completely within the 15 min of our experiment, these observations strongly suggest that pausing transcription at an early stage does not abolish loop formation or long range contacts and coordinated motion of RNA pol II bound chromatin domains.

## Conclusions

By using Optical Flow, we detected coordinated domains of chromatin motion in living human cells with nano-scale sensitivity. Chromatin domains of coherent motion exhibit long range correlation over the entire nucleus. In the absence of transcriptional activity, chromatin movements were no longer correlated independently of their constant, but small correlation length. Because protein and mRNA concentration as well as transient contacts and looping change during transcription, they also affect the mechanical properties of the chromatin fiber (20, 44, 45).

Notably, direction and magnitude of correlation length of DNA significantly differed from H2B-GFP. Despite absence of apparent defects in cell proliferation, we cannot exclude that the SiR Hoechst dye alters chromatin diffusive behavior, nor can we ascertain that the extra bulk imposed on the nucleosomes by incorporating H2B fused to GFP (a 12% or 25% increase in molecular weight of the octamer for homo- and heterotypic nucleosomes respectively) affects the analysis. Numerous previous studies suggest that the consequences should be negligible (6, 23, 24). We can, however, explain the differences in our results by a difference in labeling density and preferential incorporation of H2B-GFP into more open region of chromatin. Hence, analyzing DNA provides a more general, possibly more precise, picture with greater amplitudes between different chromatin states including inactive, dense chromatin domains, where H2B-GFP mostly informs on the behavior of more accessible chromatin.

We further show that transcription dependent motion is characterized by the appearance of vortex-like movements which are suggestive of nodes. Nodes are formed by accumulation of proteins and enzymes involved in a specific process, for example polycomb or chromatin remodelers regulating transcription (45–48). Enhancer-promoter looping or active pulling of DNA may result in vortex-like apparent motion at short time scales between domains.

The smoothness parameter allows quantification of transitions in motion and provides insight into the origin of chromatin domains. Smoothness values are characteristic of transcription induced variations in chromatin compactness. The smoothness parameter can be interpreted as a measure of smooth or sharp transitions between adjacent regions of coherent flow and therefore provides insight into time dependent formation of dynamic regions and their boundaries.

Chromatin conformation capture analysis have identified A and B compartment regrouping <10Mb domains of similar chromatin marks and compaction (3, 49). Blocks of several A or B compartments tend to interact but their assembly is stochastic and their boundaries cannot be assessed by population averaging Hi-C methods. Domains likely result from auto assembly (50) or phase separation-type physical processes driven by accumulation of proteins (51), such as for example RNA pol II factories (52), HP1 droplets (53) or repeated elements (54). On the single cell level, sharp boundaries of compartments were also seen in snapshots using super-resolution microscopy (55). Hence, the rough domain boundaries observed in this study are reminiscent of phase transitions, as they separate domains of different dynamic behavior depending on transcriptional activity. Our approach allows seeing functional domains in nuclei of living cells in real time.

## Materials and Methods

### Cell Culture

A stable human osteosarcoma U2OS cell line (ATCC) stably expressing H2B-GFP was a gift from Sébastien Huet (Rennes, France). Cells were grown in Dulbecco’s modified Eagle’s medium (DMEM) containing phenol red-free (Sigma-Aldrich) supplemented with 10% Fetal bovine serum (FBS), Glutamax containing 50 μg/ml gentamicin (Sigma-Aldrich), 1 mM sodium pyruvate (Sigma-Aldrich) and G418 0.5 mg/ml (Sigma-Aldrich) at 37°C with 5% CO2. Cells were plated for 24 h on 35 mm petri dishes with a #1.5 coverslip like bottom (μ-Dish, Ibidi, Biovalley) with a density of 100000 cells/dish.

### DNA staining

For DNA staining, the same cell line of U2OS was labeled by using SiR-DNA (SiR-Hoechst) kit (Spirochrome AG). SiR-DNA is a far-red fluorophore that binds to the DNA minor groove with high specificity (56). Briefly, 1 mM stock solution was prepared by dissolving the content of the vial of SiR-DNA in 50 μl of anhydrous DMSO. This solution should be stored at −20°C. For labeling, we diluted the stock solution in cell culture medium to concentration of 2 μM and vortex briefly. On the day of the imaging, the culture medium was changed to medium containing SiR-fluorophores and incubated at 37°C for 30-60 minutes. Before imaging, the medium was replaced by L-15 medium (Liebovitz’s, Gibco). Cells were mounted on the microscope for live imaging in a custom-built 37°C microscope incubator.

### Cell starvation, stimulation and chemical treatment

#### Transcription inhibition and stimulation

For cell starvation, the media were replaced with serumfree medium (DMEM, Glutamax containing 50 μg/ml gentamicin, 1 mM sodium pyruvate, and G418 0. 5 mg/ml). The cells were incubated for 24 h in the 37°C incubator before imaging. Just before imaging, the medium was changed to L-15 medium. Cell starvation conditions were used for transcription inhibition mode. While for stimulation mode, cells were incubated with full medium containing 10% FBS, and imaged with 10% FBS in L-15 medium.

#### Transcription blocking

To assess the impact of transcription initiation on chromatin motion in living cells, we added fresh L-15 medium containing 1 μM Triptolide (TPL, Sigma-Aldrich). To block the transcription elongation, 100 μM of 5,6-dichloro-1-β-D-ribofuranosylbenzimidazole (DRB, Sigma-Aldrich) was diluted in fresh L-15 medium and incubated under the microscope for 15 minutes before imaging.

### Microscopy and Image Acquisition

#### SiR–Hoechst labeled DNA imaging

DNA images were acquired using a DMI8 inverted automated microscope (Leica Microsystems) featuring a confocal spinning disk unit (CSU-X1-M1N, Yokogawa). Integrated laser engine (ILE 400, Andor) with a selected wavelength of 647 nm (140mW) was used for the excitation. Samples were imaged with an oil immersion objective (Leica HCX-PL-APO 100x/1.4 NA). Fluorescence emission of the SiR–Hoechst was filtered by a single-band bandpass filter (FF01-650/13-25, Semrock, Inc.). Image series of 150 frames (5 fps) were acquired using Metamorph software (Molecular Devices), and detected using sCMOS cameras (ORCA-Flash4.0 V2) and (1×1 binning), with sample pixel size of 65 nm. All series were recorded at 37°C and in a humid chamber by controlling the temperature and CO2 control flow using H201- couple with temperature and CO_2_ units.

#### H2B-GFP imaging

Series of 150 frames were acquired with an exposure time of 200 ms using a Nipkow-disk confocal system (Revolution, Andor) featuring a confocal spinning disk unit (CSU22, Yokogawa). A diode-pumped solid-state laser with single laser line was used for excitation of GFP at 488 nm (25mW; Coherent). Samples were imaged with an oil immersion objective (100X, Plan Apo 1.42, Nikon) followed by 2x magnification and fluorescence was filtered with an emission filter (ET525/30-25, Semrock, Inc.). The fluorescent emission was detected on a cooled electron multiplying charge-coupled device camera (iXon Ultra 888), with sample pixel size of 88 nm. The system was controlled using the Revolution IQ software (Andor).

#### Quantitative evaluation

A vector is determined by its direction and magnitude. The Angular Error (AE) and Endpoint Error (EE) are common measures used for performance evaluation of flow estimation methods. The Angular Error is the angle between the ground-truth vector 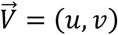 and the corresponding estimated vector 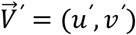 and is computed as the inverse cosine of the normalized dot-product of 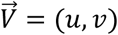 and 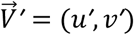 (57):

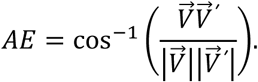

As an angle between an arbitrary vector and the zero vector is not defined, we only take non-zero displacements into consideration.

The angular error is a relative measure and penalizes discrepancies in the direction of ground-truth and estimated flow. The evaluation of errors in the magnitude is given by the absolute Endpoint Error (EE) (58).

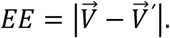

All calculations were carried out by using MATLAB (MATLAB Release 2017a, The MathWorks, Inc., Natick, Massachusetts, United States) on a 64-bit Intel Xeon CPUE5-2609 1.90 GHz workstation with 64 GB RAM and running Microsoft Windows 10 Professional.

### Image processing and data analysis

#### Denoising

Raw images are denoised using non-iterative bilateral filtering (59). While Gaussian blurring only accounts for the spatial distance of a pixel and its neighborhood, bilateral filtering additionally takes the difference in intensity values into account and is therefore an edge-preserving method. Abrupt transitions from high- to low-intensity regions (e.g. heterochromatin to euchromatin) are not over-smoothed.

#### Drift registration

Drift during image acquisition is determined by the cross correlation of the first image of the whole nucleus in the sequence and every following image. The position of the correlation peak is found with sub-pixel accuracy by a Gaussian approximation of the correlation peak. The distance of the correlation peak from the origin is the desired drift vector. The detected drift in all processed image sequences is in the range of less than 10 nm and is therefore negligible.

#### Spatial correlation calculation

The spatial autocorrelation function *r* of a scalar field γ(x,y) can be calculated efficiently by the use of Fast Fourier Transform algorithms and is given by (60):

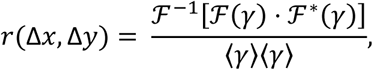

where 𝓕(·),𝓕(·),^−1^, and 𝓕*(·) are the Fourier transformation, *Inverse* Fourier transformation and the complex conjugate of the Fourier transformation, respectively. The two-dimensional autocorrelation function was calculated for horizontal and vertical space lag as denoted by *r*(Δ*x*, Δ*y*). One can project the two-dimensional correlation function onto one dimension using the space lag ρ^2^ = Δ*x*^2^ + Δ*y*^2^. The projection was carried out as a radial average and the correlation function becomes a function of the space lag only, i.e. *r* = *r*(*ρ*).

## Acknowledgements

We thank Sébastien Huet (IGDR-UMR6290, Rennes, France) for providing the U2OS H2B-GFP cell line. Alain Kamgoue helped with setting up the Amazon EC2 computation cluster that was used for our calculations. We acknowledge support from the TRI imaging platform, Toulouse. KB was supported by the ANR ANDY, IDEX ATs NudGENE and the Foundation ARC.

## Supporting Information

Supplementary Note 1 contains details on fundamental functionality and limitations of tested Optical Flow methods, simulated microscopic and dynamic parameters along with further analysis of the accuracy between methods. Supplementary Note 2 contains details on of the correlation on circular variables, correlation models and the smoothness parameter.

## Author contributions

H.A.S. and K. B. designed the research; H.A.S. performed all experimental work; R.B. implemented and developed the data processing algorithm; H.A.S. and R.B. analyzed the data; H.A.S. supervised R.B.; K. B. funded the research; H.A.S., R. B. and K. B. wrote and edited the manuscript.

